# Cep57 is a cohesin-associated regulator of chromosome segregation, cell cycle progression and genome stability in early embryos

**DOI:** 10.1101/2025.04.10.648303

**Authors:** Sharada Iyer, Lakshmi Prasanna Sai Madamanchi, Advait Gokhale, Megha Kumar

## Abstract

Faithful chromosome segregation and coordinated cell-cycle progression are essential for maintaining genome stability and preserving progenitor populations during vertebrate embryogenesis. Although pathogenic CEP57 variants are associated with developmental disorders including aneuploidy and microcephaly, the molecular mechanisms by which CEP57 coordinates these processes remain poorly understood. Here, we identify Cep57 as a previously unrecognized regulator of chromosome segregation, G1/S transition, centrosome integrity, and genome surveillance during early zebrafish embryogenesis. Cep57 localizes to both the nucleus and centrosomes of zebrafish blastulae, indicating probable dual functions in nuclear cell cycle regulation and centrosome organization. Our results show that Cep57 associates with Rad21, Smc1, and Smc3, revealing an unexpected functional link between Cep57 and the cohesin complex. Loss of Cep57 results in Rad21 depletion, destabilization of the cohesin complex, chromosome segregation errors resulting in supernumerary nuclei, and disruption of pericentriolar material organization, demonstrating a critical role in maintaining mitotic fidelity. We further show that Cep57 interacts with Geminin and promotes an Rb1-dependent G1/S checkpoint, whereas Cep57 deficiency causes widespread cell cycle dysregulation, genome instability, and apoptosis. Quantitative proteomic analyses reveal robust activation of DNA damage and checkpoint signaling pathways, consistent with engagement of genome surveillance mechanisms following Cep57 loss. Importantly, these cellular abnormalities precede extensive neural apoptosis, depletion of neuroprogenitor populations, and the emergence of microcephaly-associated phenotypes. Collectively, our findings establish Cep57 as a cohesin-associated integrator of chromosome segregation, cell cycle progression, and genome stability that is essential for neural progenitor maintenance during vertebrate embryogenesis, thereby expanding its functional repertoire far beyond its canonical role in centrosome regulation.

## Introduction

Centrosomal protein 57 (CEP57) plays key roles in centriole dynamics, mitotic spindle formation, and organization. It is essential for pericentriolar matrix (PCM) organization, centriole duplication, and engagement. Lack of CEP57 results in PCM disorganization and precocious centriole disengagement, resulting in ectopic Microtubule Organizing Centres (MTOCs).^1–7^ It also plays key roles in the nucleation and stabilization of microtubules and spindle formation, and its depletion results in misaligned chromosomes and multipolar spindles.^4,8,9,10^ Further, it localizes to the kinetochores and interacts with microtubules to remove MAD1, and its abrogation results in impaired Spindle Assembly Checkpoint (SAC) signaling and chromosome segregation defects.^7,9,11^ CEP57 also localizes to the cytokinetic midbody and aids in central spindle microtubule organization.^12^ Mutations in the *CEP57* gene are associated with Mosaic Variegated Aneuploidy syndrome (MVAS). MVAS is an autosomal recessive disorder characterized by chromosome gain or loss in somatic cells, which results in mosaic aneuploidies. These patients show microcephaly, skull deformities, facial dysmorphism, and predisposition to tumors. ^1,6,13–15^ The *Cep57* homozygous knockout was embryonic lethal, and the heterozygous mice exhibited severe vertebrate ossification defects and died shortly after birth. The cells showed aneuploidy, impaired centrosomal maturation, premature centriole disjunction, and chromosome segregation anomalies.^4,16^ However, the mechanistic underpinnings of microcephaly mutations in *CEP57* remain unknown. Recent studies implicate the tight interplay of centrosomal pathways with DNA damage response to maintain genome integrity.^17^ Yet, how centrosomal defects are communicated to cell cycle checkpoints during embryogenesis is poorly understood. Hence, understanding the contribution of centrosomal dysfunction to developmental disorders remains elusive.

As CEP57 has been classically studied in centrosome-dependent mitotic events, and CEP57-associated MVA has been attributed to mitotic anomalies, our study expands its functional repertoire. Here, we explored non-centrosomal functions of Cep57, its role in cell cycle progression and cell fate specification during early embryogenesis (Fig. 1A). We demonstrate that Cep57 interacts with cohesin complex members – Rad21, Smc1 and Smc3 to maintain genomic integrity and cell cycle progression during early embryonic development. We show that Cep57 stabilizes the cohesin complex and ensures faithful chromosome segregation during early cell divisions. The Cep57-depleted embryos exhibit segregation errors, supernumerary micronuclei reflecting aneuploidy, genome instability, and impaired cell cycle progression. Our experiments reveal a previously unrecognised regulatory interaction between Cep57 and an important G1/S checkpoint regulator, Geminin. Loss of Cep57 results in elevated levels of Geminin and, consequently, disruption of G1/S progression, culminating in G1 arrest. This leads to extensive apoptosis in the embryo, along with neurodevelopmental defects, resulting in microcephaly-associated phenotypes. To summarise, in addition to Cep57’s role in centrosome-associated mitotic errors, we identify Cep57 functions in G1/S progression and safeguarding genome stability in early embryogenesis. These cellular defects are coupled to neurodevelopmental anomalies, increased neural tissue apoptosis, reduced expression of neural crest specification genes, resulting in microcephaly-associated features.

**Figure 1:**
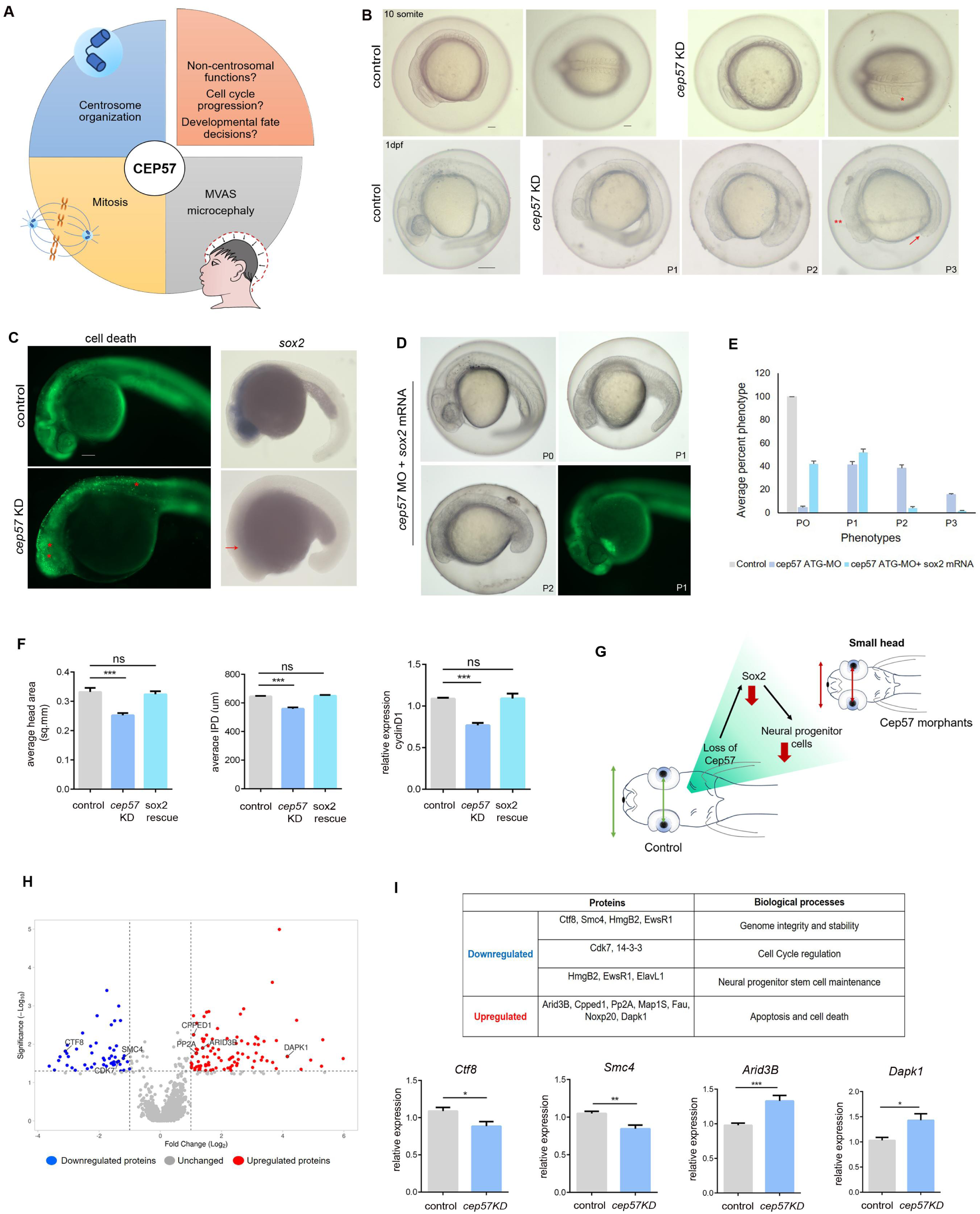
Cep57 localizes to centrosomes and is essential for early neurodevelopment. A: Sum projection confocal images of 256 cell stage WT embryos showing centrosomal staining by γ tubulin (red), Cep57 (green, red asterisks), and DNA (blue/DAPI). Scale bars, 50µm. B: Western blot showing localization of Cep57 in whole embryonic lysate, nuclear and cytoplasmic fractions. Lamin B1 and Gapdh represents nuclear and cytoplasmic fractions respectively. C: Western blot showing Cep57 levels in control and morpholino-injected embryos. Gapdh was used as the loading control. D: Quantification showing percent phenotype of *cep57* splice morpholino injected, splice morpholino coinjected with *cep57* mRNA, and *cep57* mRNA only. Data are shown as mean ± SEM, *p<0.05, **p<0.01, ***p<0.001. n=3 for each experiment. E: Gross morphological analysis of *cep57-*depleted embryos in comparison to control at 10 somite and 24hpf stage, showing somite architecture anomalies (red asterisk), midbrain-hindbrain boundary (double red asterisk), and body axis defects (red arrow). Scale bars, 100µm for 10 somite and 50µm for 24hpf embryos. F: Whole mount skeletal preparations of 5dpf control and *cep57* morphant larvae showing cartilage and bone staining (red asterisk) using Alcian blue and Alizarin red S, respectively. Scale bars, 100µm. G: Quantification of microcephaly-associated features – interpupillary distance (IPD) and head area in control and *cep57* morphants. Data are shown as mean ± SEM, *p<0.05, **p<0.01, ***p<0.001. n=3 for each experiment.

## Results

### Cep57 is essential for early neurodevelopment and maintenance of head size

To delineate the developmental roles of *cep57*, we first examined its spatio-temporal expression across developmental stages. *cep57* was expressed ubiquitously during cleavage stages (2-cell stage), suggesting that it is maternally inherited and continued to be expressed till the somitogenesis phase (Fig. S1A, B). The spatio-temporal expression of *cep57* was restricted to the eye and brain during the later phase of somitogenesis until 1dpf /24hpf (red asterisk, Fig. S1A). Cep57 protein levels were also maintained across early developmental stages till 1dpf (Fig. S1C). To ascertain its developmental functions, we used a gene knockdown approach with sequence-specific antisense morpholinos (translation and splice-site blockers) against zebrafish *cep57*.

The *cep57* translation blocker morphants showed abnormalities in the brain architecture and organization, eye and otic vesicle defects (double red asterisks, Fig. 1B) somitic defects (red asterisk, Fig. 1B), along with a poorly developed anteroposterior axis (red arrow, Fig. 1B). We categorized the morphants as normal, similar to controls (P0, not shown), embryos with brain abnormalities (P1), embryos with brain abnormalities and anterior-posterior axis defects (P2), and the extremely severe phenotype with poorly developed brain and body axis (P3) (Fig. 1B). The severity of the phenotypes P0, P1, P2, and P3 increased in a morpholino concentration-dependent manner (Fig. S1D). Cep57 depletion was confirmed by immunoblotting with the morphant lysates (Fig. S1E). To ascertain the specificity of the morpholinos and the resulting phenotype, we performed rescue experiments by co-injecting the splice-blocker morpholino and zebrafish *cep57* mRNA into 1-cell-stage embryos. The co-injection of *cep57* mRNA significantly rescued all the phenotypes (Fig. S2A, B). Thus, *cep57* is crucial for neurodevelopment and axis specification during early embryogenesis.

To determine the cellular basis of the poorly developed brain, we performed a cell death assay. The embryos showed extensive apoptosis/cell death, marked by acridine orange-positive cells in the brain and spinal cord (red asterisks, Fig. 1C) and fluorescein-based terminal deoxynucleotidyl transferase dUTP nick end labelling (TUNEL) staining (white asterisks, Fig. S2C). During early neurodevelopment, *sox2* is essential for the proliferation and maintenance of neural progenitors.^20,22,23^ Interestingly, we observed a severe reduction of *sox2*-positive cells in the developing CNS, suggesting a severe loss of the neuroprogenitor population upon *cep57* depletion (red arrow, Fig. 1C). The severe phenotypes P1, P2, and P3 and cell death were significantly rescued in cep57 morphants co-injected with zebrafish *sox2* mRNA (Fig. 1D, E).

At larval stages, the morphants showed gross craniofacial defects, cardiac edema (red asterisk), and severe anteroposterior axis defects (red arrowhead) (Fig. S2D). The P1 and P2 morphants showed severe craniofacial dysmorphic features, cartilage, and skeletal abnormalities at larval stages (double red asterisk, Fig. S3A). The morphant larvae exhibited reduced head size and interpupillary distance (IPD) (Fig. 1F), strongly indicative of a microcephaly-like phenotype. To confirm the microcephaly-like phenotype, we also examined the levels of microcephaly-associated genes *mcph1, wdr62, ankle2, map11, kif14*, and *aspm* (Fig. S4). The *cep57* morphants exhibited a marked reduction in the expression of all these microcephaly-associated genes (Fig. S4). The reduced expression of microcephaly markers indicates that cep57 plays a key role in maintaining neural progenitor cell fate and regulating head size. This decrease in head area, IPD, and craniofacial anomalies was significantly reduced upon *sox2* rescue (Fig. 1F, S3B). Further, introduction of *sox2* also restores *cyclin D1* levels and hence promotes proliferation in *cep57* morphants (Fig. 1F). These results show that the maintenance of undifferentiated progenitor fate and proliferation is regulated by *cep57*-mediated *sox2* activation. The reduced sox2 expression in *cep57* morphants leads to loss of neuroprogenitor cells and results in microcephaly. (Fig. 1G)

The craniofacial defects can be attributed to the reduced expression of neural crest specifier genes like *crestin* and *sox10*, as the craniofacial skeletal elements primarily develop from neural crest-derived mesenchyme.^18–20^ We observed that *cep57* depletion resulted in the reduced expression of cranial neural crest specifiers *crestin*, *snail2* and *sox10* (Fig. S5). *crestin* is a pan-neural crest marker, expressed in premigratory and migrating neural crest cells.^21^ *snail2* and *sox10* are expressed in the premigratory cranial neural crest cells and are required for neural crest survival, specification, and migration.^20^ We observed severe down-regulation of *crestin*, *snail2,* and *sox10* in the *cep57* morphants at the early somitogenesis phase (red asterisk, Fig. S5A, B, C) and 24hpf P2 morphants (double red asterisk, Fig. S5D, E, F). The down-regulation of these specifier genes strongly suggests that *cep57* is essential to maintain the neural crest progenitor fate. Hence, gene expression analysis highlights the role of *cep57* in cranial neural crest specification and mesoderm patterning.

To probe further into the mechanistic basis of the *cep57* knockdown phenotypes, we performed label-free LC-MS/MS proteomics analysis to identify differentially regulated proteins in *cep57* morphants (Fig. 1H). We confidently identified 121 upregulated and 65 downregulated proteins in the *cep57* morphants (Fig. S6). Owing to the limited mapping of the zebrafish proteome in multiple databases such as Panther, String, and ShinyGo, a total of 19 GO terms were identified, including biological processes for the upregulated category and a total of 18 GO terms, including molecular function, for the downregulated proteins (Fig. S6). The GO terms of the fold-enriched biological processes were apoptosis, microtubule dynamics, vesicular trafficking, and cytoskeletal organization in the upregulated protein category (Fig. 1H, S6). The analysis revealed upregulation of pro-apoptotic proteins such as Arid3B, Cpped1, Protein phosphatase 2A (Pp2A), and Death-associated protein Kinase 1 (Dapk1) (Fig.1H, I). We also observed a corresponding upregulation of *dapk1* and *arid3b* transcripts in the morphants (Fig. 1I). ARID3B activates pro-apoptotic p53 target genes and induces apoptosis.^37^ The overexpression of CPPED1 results in impaired cell cycle progression and promotes apoptosis.^38^ PP2A upregulation leads to caspase 3 activation and hence has a pro-apoptotic function.^39^ DAPK1 is a key regulator of cell death and mediates caspase-dependent and caspase-independent pro-apoptotic signaling. Recent reports also show its function in tumor suppression and neuronal cell death.^40–42^ Thus, proteome profiling strongly suggests that Cep57 is essential for cell survival during early embryogenesis.

For downregulated proteins, the GO terms for fold-enriched biological processes were chromosome condensation, sister chromatid cohesion, and segregation coupled with cell cycle regulators (Fig. 1H, I, S6). The quantitative proteome revealed downregulation of Ctf8, Smc4, 14-3-3, Cdk7, and neuroprogenitor maintenance factors such as ElavL1 (Fig. 1H, I). CTF8 is essential for the establishment of sister chromatid cohesion, and its depletion results in cohesion defects, chromosome segregation defects, and cell cycle arrest.^45,46^ SMC4 is part of the condensin complex, and its phosphorylation induces G1 arrest and plays a key role in single-strand DNA break repair.^47–49^ The downregulation of Ctf8 and Smc4 indicate the role of Cep57 in genome survellience functions. 14-3-3 proteins function at G1/S transitions and regulate the timing of mitosis.^43,44^ In cycling cells, CDK7 phosphorylates CDK4 and CDK6 to initiate G1 progression and CDK2 for G1/S transition. The G2/M transition is also induced by CDK7-mediated phosphorylation of CDK1, and inhibition of CDK7 results in BAK and effector caspase activation.^51–53^ Mice lacking CDK7 showed increased apoptosis, and it is essential for neural progenitor proliferation.^50^ The downregulation of Cdk7 and neuroprogenitor maintenance genes corroborates with increased apoptosis and loss of sox2-positive neuroprogenitor cells (Fig. 1C) in *cep57*-depleted embryos. Together, quantitative mass spectrometry-based proteome profiling reinforces that loss of *cep57* results in increased cell death, loss of neuroprogenitor cells coupled with impaired cell cycle progression (Fig. 1).

### Cep57 is essential for cell cycle progression

To examine if *cep57* plays a role in cell cycle progression during early development, we performed fluorescence-based cell cycle analysis (Fig. 2A). Remarkably, we observed a significant increase in the G1 phase and concomitant reduction in the G2/M phase in the 1dpf *cep57* morphants when compared to control. The S phase remained unchanged in the morphants (Fig. 2B, D). As development proceeded in time by 2dpf, the accumulation of cells in the G1 phase resulted in a consequent reduction in the S phase, showing impaired G1/S transition (Fig. 2C, D). The whole embryo DNA content flow cytometry assay was validated by immunoblotting for cell cycle markers (Fig. 3E). Geminin is one of the master regulators of cell cycle progression, accumulating in S, G2, and M phases.^25^ At G1, Geminin levels are low; during G1-S transition, Geminin accumulates to initiate DNA replication. High levels of Geminin in the G2 and M phases ensure that DNA re-replication is inhibited. At metaphase-anaphase transition, Geminin is degraded to prepare the cell for the next cycle. ^26,27^ Ectopic expression of Geminin in G1 elicits cell cycle arrest, failure to progress into S phase, and apoptosis. ^28^ During cleavage (64-128 cell stage), Geminin levels are unchanged in the *cep57* morphants (Fig. 3E). However, Geminin levels increased as development proceeds in time, particularly in the sphere stage, gastrulation (50% epiboly) until 1dpf, suggesting that Cep57 depletion resulted in G1 arrest (Fig. 3E).

**Figure 2:**
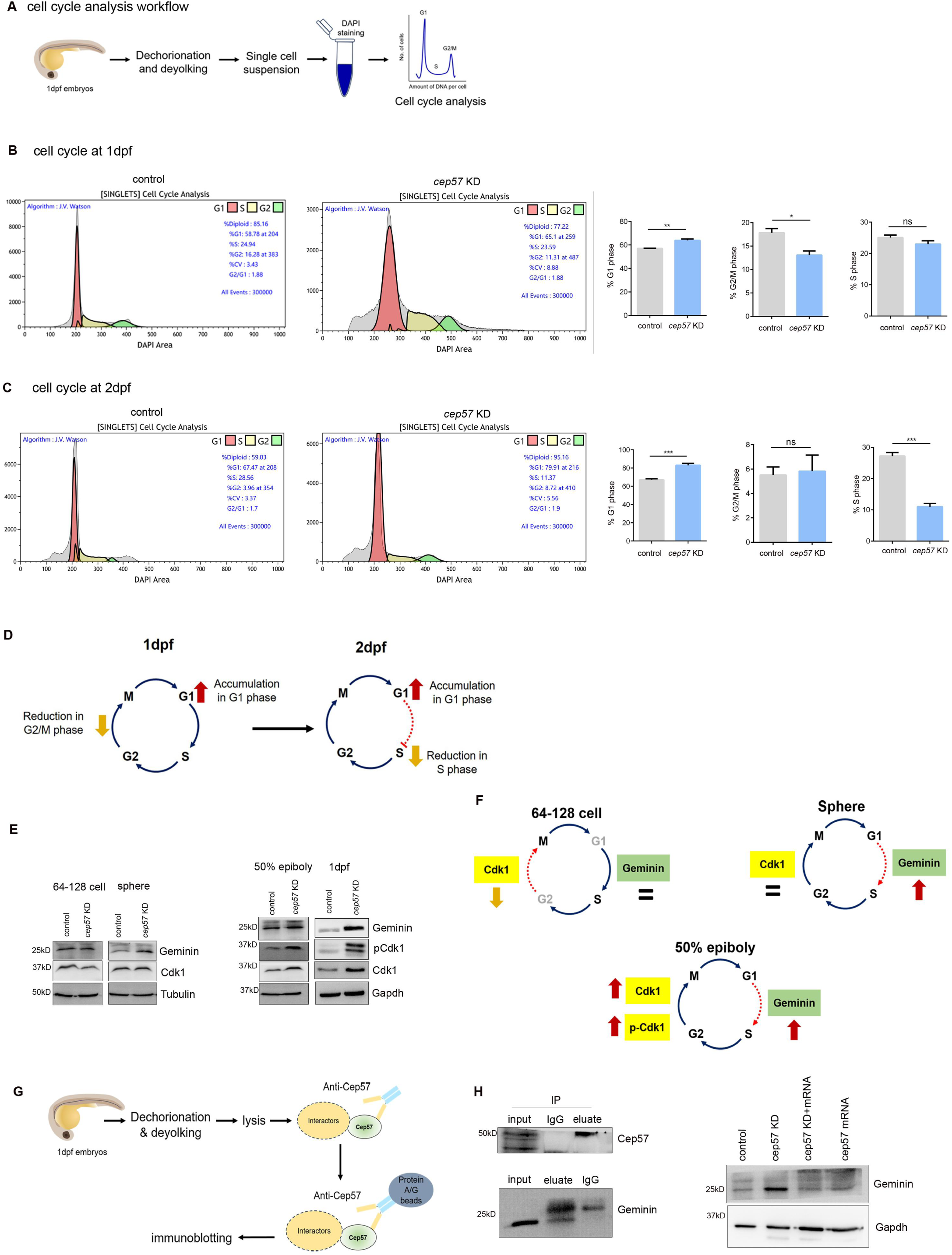
Cep57 is essential for cell survival, genome stability and cell cycle progression. A: Whole-mount acridine orange staining to visualize apoptotic cells (red asterisks) in control and *cep57* morphants. Scale bar, 100µm. B: Whole-mount TUNEL staining in control and *cep57* morphants. Scale bar, 100µm. C, D: Fluorescence-based flow cytometry analysis in control and *cep57* morphants showing different phases of the cell cycle at 1dpf (C) and 2dpf (D). E, F: Quantification of cells in each phase of the cell cycle in control and cep57 morphants at 1dpf (E) and 2dpf (F). Data are shown as mean ± SEM, *p<0.05, **p<0.01, ***p<0.001. n=3 for each experiment.

**Figure 3:**
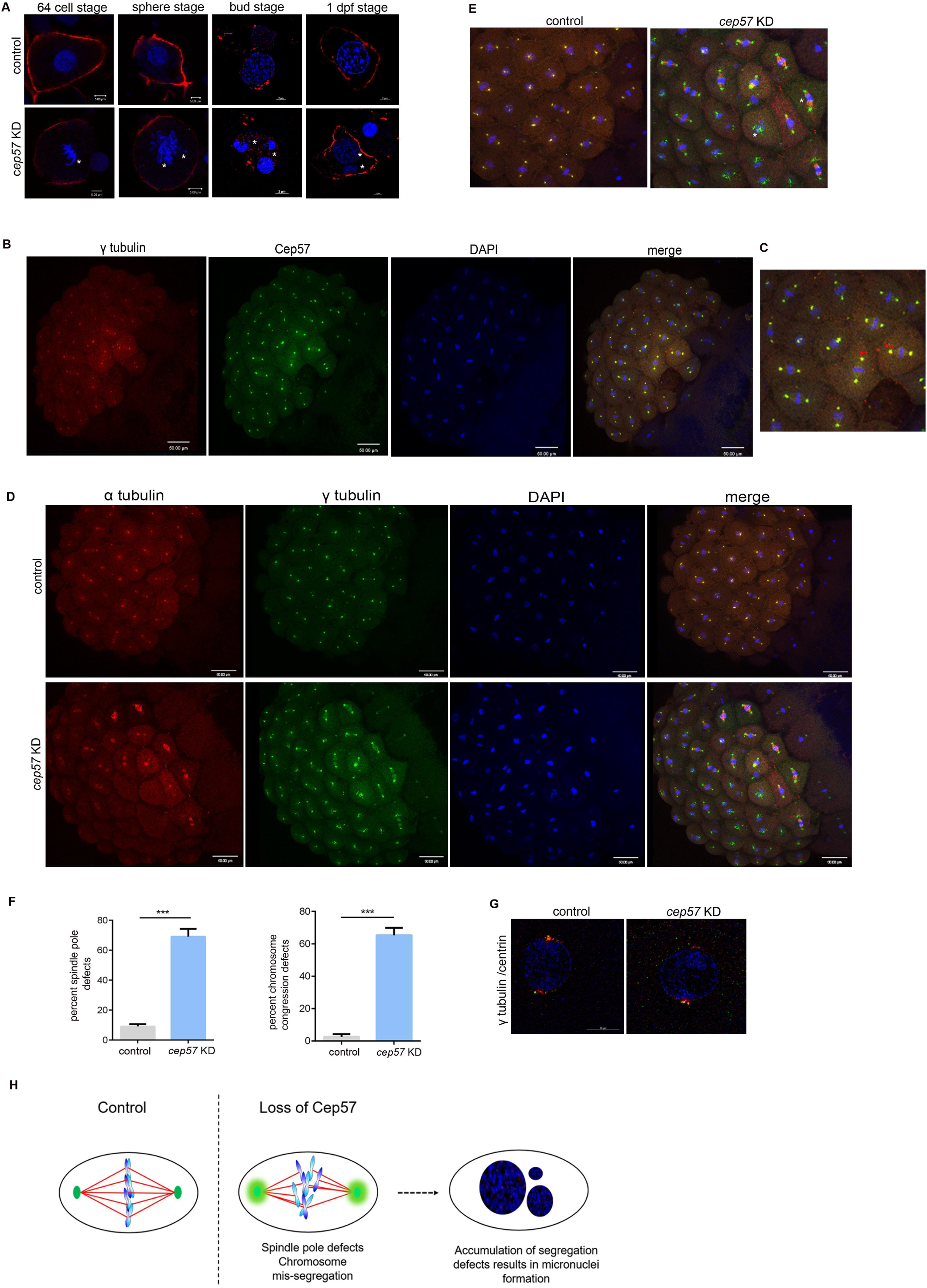
Loss of Cep57 results in PCM disorganization, DNA damage, and supernumerary micronuclei. A: Western blot showing Geminin, Cdk1 and phospho Cdk1 levels in control and morpholino-injected embryos across developmental stages. Gapdh or Tubulin was used as the loading control. B, C: Sum projection confocal images of 256 cell stage control and *cep57* morphant embryos showing microtubule and centrosomal staining at metaphase (B) and interphase (C) by α tubulin (red), γ tubulin (green, red asterisks) respectively, and DNA (blue/DAPI). Scale bars, 50µm. D: Quantification of *cep57* morphant embryos showing spindle pole focusing defects. All data are shown as mean ± SEM. *p<0.05, **p<0.01, ***p<0.001. n=3 for each experiment. E: Sum projection SIM^2^ images of 256 cell stage control and *cep57* morphants showing centrosomal staining γ tubulin (red), centrioles (green) and DNA (blue/ DAPI). Scale bars, 10µm. F: Sum projection SIM^2^ images of cells from control and *cep57* morphants showing cell membrane (actin red) and DNA (blue/ DAPI) at bud stage and 24hpf. Scale bars, 2µm.

The progression throughout the cell cycle also requires the regulatory oscillatory activity of cyclins and cyclin-dependent kinases (Cdks). Cyclin-dependent kinase 1 (Cdk1) regulates G1 to S and G2 to M phases, and its activity is modulated by phosphorylation and binding to Cyclin B1.^29^ Cdk1 levels are low in G1, and activity increases by late G1 until anaphase. During cleavage stages, we observed a reduction of Cdk1 levels, corroborating the flow cytometry analysis of reduced G2/M entry upon cep57 depletion (Fig. 3E, F). The Cdk1 levels remain unchanged until the sphere stage in *cep57* morphants (Fig. 3E, F). Interestingly, post-gastrulation (50% epiboly), we observed a concomitant increase in the inhibitory form of Cdk1, phosphor-Cdk1 (Tyr15), as well, suggesting that Cdk1 is maintained in the inhibitory state upon *cep57* depletion, strongly indicating inhibition of cell cycle progression and G1 arrest (Fig. 3E, F).

To determine molecular regulators of cell cycle progression in *cep57* morphants, we performed endogenous immunoprecipitation assays to identify Cep57-mediated G1/S checkpoint regulators. (Fig. 2G). Our endogenous immunoprecipitation experiments showed that Cep57 interacts with Geminin (Fig. 2H), suggesting that the Cep57-Geminin interaction is essential for the G1/S transition. The loss of Cep57 results in Geminin accumulation, leading to G1 arrest. Indeed, the overexpression of Cep57 mRNA restored the Geminin levels in the Cep57-depleted embryos, allowing normal cell cycle progression (Fig. 4H).

**Figure 4:**
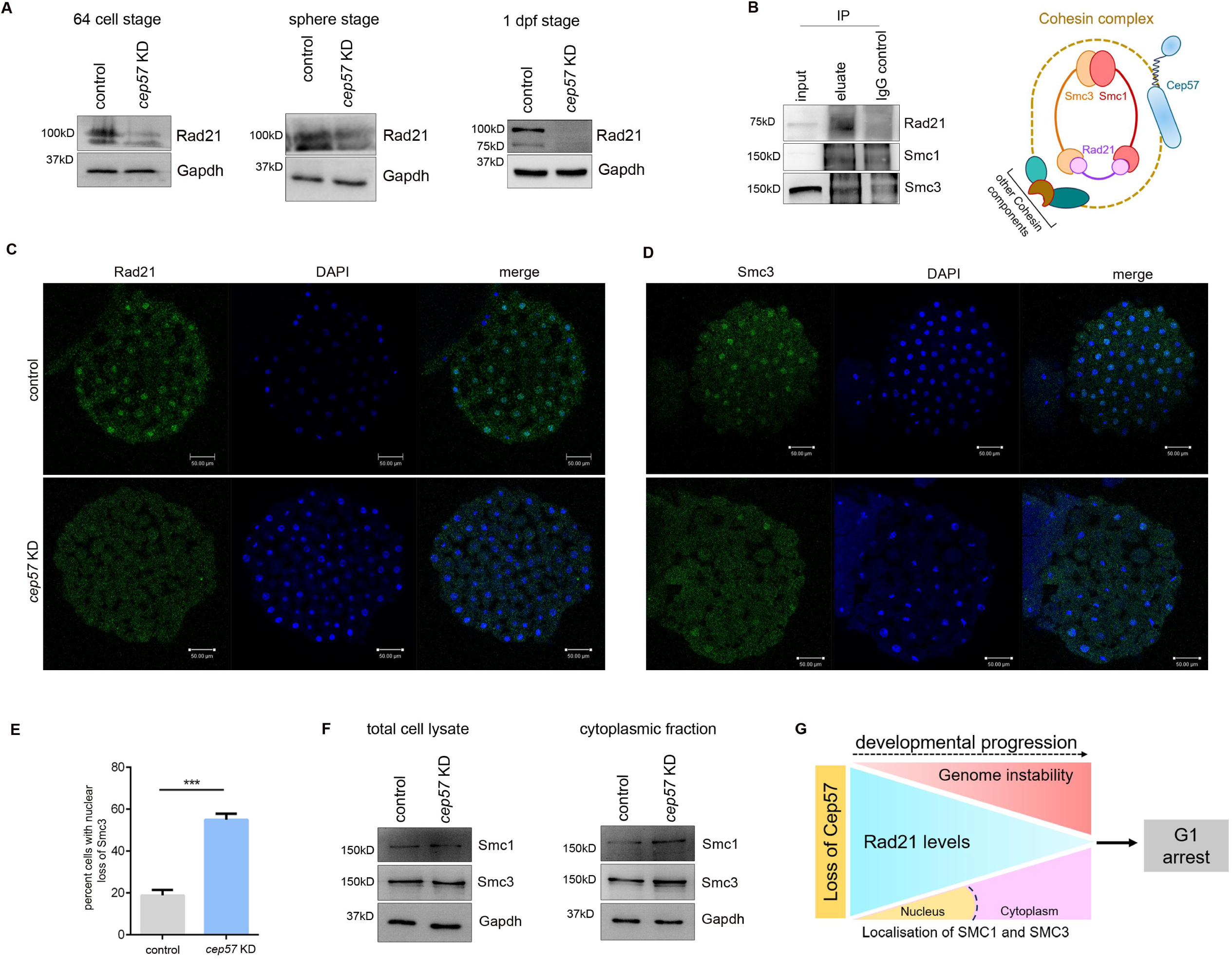
Cep57 interacts with Geminin to regulate Rb1-mediated G1/S transition. A: Comet assay to estimate DNA damage in control and *cep57* morphants. All data are shown as mean ± SEM. *p<0.05, **p<0.01, ***p<0.001. n=3 for each experiment. B: Western blot showing Rad21 levels in control and morpholino-injected embryos. Gapdh was used as the loading control. C: Immunoblot showing Rad21 as an interactor of Cep57 in endogenous immunoprecipitation assay. D: Quantitative real-time PCR plots showing relative transcript levels of *p53* and *p21*. All data are shown as mean ± SEM. *p<0.05, **p<0.01, ***p<0.001. n=3 for each experiment. E: Western blot showing Rb and phospho Rb levels in control and morpholino-injected embryos. Gapdh was used as the loading control. F: Quantification of Fluorescence-based flow cytometry analysis in control and *cep57* morphants showing different phases of the cell cycle in control, cep57 knockdown and Rb mutant background. Rb mutant background served as positive control. All data are shown as mean ± SEM. *p<0.05, **p<0.01, ***p<0.001. n=3 for each experiment. G: Western blot showing Geminin levels in control, *cep57* morphants and *cep57* depletion in Rb mutant background. Gapdh was used as the loading control. H: Immunoblot showing Geminin as an interactor of Cep57 in endogenous immunoprecipitation assay. I: Western blot showing Geminin levels in control, *cep57*-depleted embryos, and *cep57* depleted embryos injected with Cep57 mRNA. Gapdh was used as the loading control.

### Loss of Cep57 results in PCM disorganization, supernumerary micronuclei, and genome instability

The impaired G1/S transition intrigued us to probe for the effect on M phase upon *cep57* depletion. We thus examined the morphants for mitotic aberrations during early embryonic divisions (Fig. 3). We observed chromosome segregation defects during cleavage divisions (64-cell stage) and post-gastrulation (sphere stage) (white asterisks, Fig. 3A). By initiation of segmentation (bud stage), we observed the presence of supernumerary micronuclei in *cep57* morphants (white asterisks, Fig. 3A), strongly indicating aneuploidy and genomic instability due to the early chromosome segregation defects. Notably, Cep57 localizes to the spindle poles in metaphase (double red asterisks, Fig. 3B, C), and in the nucleus (white asterisk, Fig. S7A, B). of zebrafish blastulae. Cep57-depleted embryos showed spindle pole focusing and pericentriolar matrix defects at midblastula transition (white asterisks, Fig. 3D, E, F), highlighting the centrosomal function of *cep57* in maintaining spindle pole integrity, in a manner akin to the *Xenopus* egg extract system.^7^ However, the centriolar organization remained unperturbed upon Cep57 depletion (Fig. 3G). Hence, Cep57 is essential to maintain spindle pole integrity, faithful chromosome segregation, and genome stability during early embryogenesis (Fig. 3H).

In order to quantify genomic instability and DNA damage in *cep57* morphants, we used the comet assay (Fig. S8A). The broken DNA fragments formed a typical “tail” in the morphant samples, showing that Cep57 depletion results in single/ double-stranded DNA breaks and the formation of alkali-labile sites (Fig. S8A). Further, we used a random amplified polymorphic DNA (RAPD) PCR assay to check for genomic instability, DNA alterations, damage, and mutations. ^31,32^ The amplification products obtained showed many bands of molecular size between 300 to 1500bp. The *cep57* morphants showed variation in band intensity as well as loss of polymorphic bands in the amplification patterns with respect to the control (red arrows, Fig. S8B). We observed the disappearance of bands in *cep57* morphants, corroborating a 38.7% reduction in genomic template stability (GTS%) (Fig. S8B). ^31,32^ Also, the DNA damage-induced G1 to S kinase, Cyclin E levels increased in the morphants, which also results in G1 arrest (Fig. S8C).

The *cep57* morphants showed reduced PH3 staining, indicating a reduction in M-phase cells (Fig. S5C)

### Cep57 interacts with the cohesin complex to maintain genome stability

Genomic instability is attributed to unequal chromosome segregation which leads to DNA damage responses. To determine the molecular basis of genomic instability in cep57 morphants, we examined Rad21, a core cohesin complex member that enables the separation of sister chromatids. The depletion of Rad21 results in impaired DNA damage response and is critical for DNA damage repair. In addition to proteolytic cleavage during mitosis, it is also cleaved during nuclear fragmentation in apoptosis.^33,34^ The morphants showed depletion of Rad21 protein across developmental stages, indicating abrogation of Rad21 mediated DNA damage response upon *cep57* depletion (Fig. 4A, 4C). Interestingly, the endogenous immunoprecipitation assays revealed that Cep57 interacts with Rad21, other cohesion complex members Smc1 and Smc3 (Fig. 4B). Cep57 depletion also resulted in loss of nuclear Smc3 (Fig. 4D, E). While the total Smc1 and Smc3 protein levels do not change upon cep57 depletion, there is nuclear loss and a concomitant enrichment of Smc1 and Smc3 in the cytoplasmic fractions (Fig. 4F). Together, we show that Cep57 interaction with cohesin members Rad21, Smc1 and Smc3 is essential for proper chromosome segregation and functioning of the DNA damage repair machinery. The loss of Cep57 results in the abrogation of Rad21 and the destabilization of the cohesin complex, resulting in chromosome segregation errors, formation of extranuclear bodies or micronuclei coupled with extensive DNA damage, culminating in G1 arrest (Fig. 4G).

### Cep57- Geminin interaction regulates Rb1-mediated G1/S transition

DNA damage also induces the activation of the cell cycle checkpoint regulatory pathway involving p53-p21-Retinoblastoma (Rb) signaling. Rb1 is a G1/S checkpoint regulator; in its active or hypophosphorylated form, it binds to the E2F transcription factor, represses target genes, preventing cells from entering S phase and propagating damaged DNA.^36^ Upon G1/S transition, when phosphorylated by CDKs, it releases E2F, resulting in cells to progress into S phase (Fig. 5A). The *cep57* morphants showed reduced levels of phospho form of Rb1, indicating that Rb1 is maintained in inactive state upon Cep57 depletion and hence resulting in inhibition of G1 to S transition (Fig. 5B). DNA damage also causes p53 activation, which in turn induces *p21* expression. The high levels of p21 result in the Rb-E2F complex formation, downregulation of cell cycle genes, and finally lead to cell cycle arrest.^35^ The *cep57* morphants showing DNA damage, also showed increased *p53* expression and consequently, high levels of *p21* expression (Fig. 5C). Next, we used Rb1 mutant zebrafish embryos to confirm if Cep57 induced G1 arrest is mediated by Rb1 (Fig. 5D). The increase in G1 phase and decrease in G2/M phase upon Cep57 depletion were rescued significantly in the Rb mutant background (Fig. 5D). Further, Geminin and *sox2* levels were also restored in the Cep57-depleted Rb1 mutant background (Fig. 5E). These findings collectively show that the loss of Cep57 results in an Rb1-mediated G1 arrest in the embryo.

**Figure 5:**
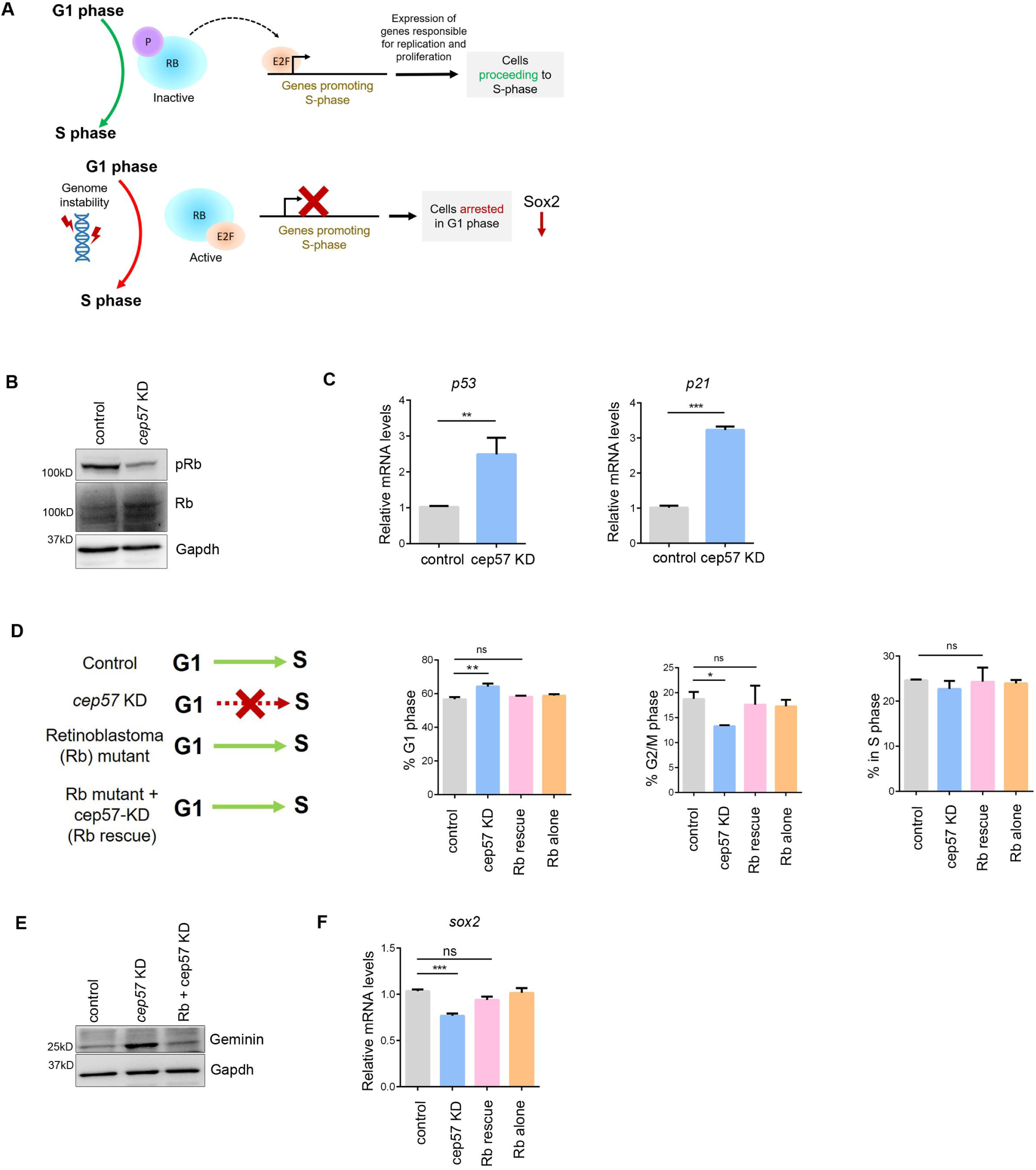
Cep57 regulates cell cycle modulators and DNA damage-inducing proteins. A: Volcano plot showing fold change in differentially expressed proteins in control vs *cep57* morphants. B: Quantitative real-time PCR plots showing relative transcript levels of *smc4, ctf8, dapk1* and *arid3b* in control and *cep57* morphants. All data are shown as mean ± SEM. *p<0.05, **p<0.01, ***p<0.001. n=3 for each experiment. C: Schematic representation showing *cep57* function in genome stability, faithful chromosome segregation, cell cycle progression, and early neurodevelopment.

## Discussion

CEP57 has classically been recognized for its pivotal role in centrosomal functions and mitotic events such as centriole biogenesis, organization of microtubule-based bipolar spindle, and faithful chromosomal segregation. Clinically, mutations in the *CEP57* gene have been linked to MVA, primarily characterized by microcephaly. At the cellular level, microcephaly is often associated with aberrations in centrosomal function, genome instability, and aneuploidy. The development of microcephaly is attributed to the failure of self-renewal or accumulated DNA damage in neural progenitor cells owing to checkpoint defects, DNA repair deficiencies, mitotic errors, and increased apoptosis. These findings underscore the importance of the interplay between the DNA damage response and centrosomal pathways in understanding the basis of microcephaly. Notably, several genes implicated in microcephaly, such as *MCPH1* and *CEP152*, possess dual roles in both DNA damage response and centrosomal function.^54^

In this study, we probe the developmental functions of *cep57* and the mechanistic basis of *cep57*-associated MVA and microcephaly. We delineate previously uncharacterized, novel molecular interactions of Cep57 with cohesin complex members - Rad21, Smc1, and Smc3 to maintain genome integrity and cell cycle progression. During early embryonic mitoses, loss of Cep57 results in destabilization of the cohesin complex, resulting in chromosome segregation errors, extra nuclear bodies (aneuploidy), and varied mitotic aberrations. After the midblastula transition, which marks the introduction of G1 phase in the cycling embryonic cells, these mitotic defects result in an Rb1-mediated G1 arrest, leading to a surge of apoptosis coupled with loss of neuroprogenitor cells, resulting in a microcephaly-like phenotype. (Fig. 6).

**Figure 6:**
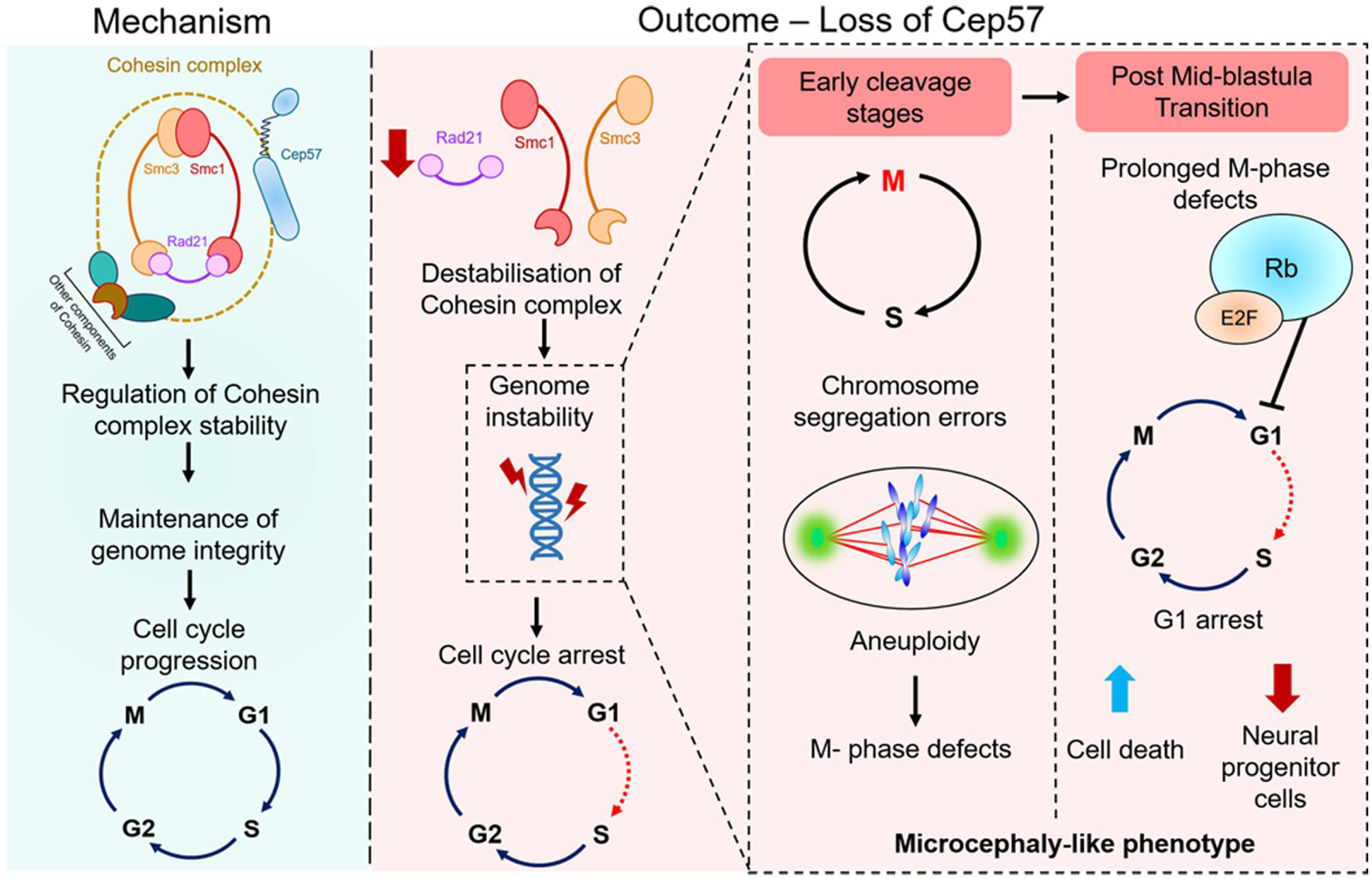

As the functions of *cep57* have largely been centrosomal, often associated with mitotic events, the MVA-associated phenotypes were attributed to these mitotic aberrations. While our findings corroborate the centrosomal mitotic anomalies, we also highlight a significant contribution of disrupted G1/S transition, DNA damage, and genome instability to the observed phenotypes. While we provide strong evidence that Cep57 is a key regulator of G1/S cell cycle progression, we cannot entirely rule out functional redundancy with other Cep family members, particularly regarding its mitotic roles, as we have investigated the cellular and developmental functions in an *in vivo* animal model-based knockdown system. Furthermore, although our proteomic analysis yielded a limited set of candidate proteins, likely due to incomplete mapping of the zebrafish proteome in public databases, it nevertheless identified several critical proteins involved in chromosome organization, cell cycle regulation, and apoptosis. In summary, we identify novel, multifaceted functions of *cep57* in cell cycle progression, maintenance of spindle pole integrity, faithful chromosome segregation, and genome stability during vertebrate embryogenesis. Our findings open new avenues for investigating broader roles of *cep57* functions in other cell cycle checkpoints, apoptotic regulation, and maintenance of chromosome integrity, which will be crucial to uncover its diverse cellular functions.

## Methods

### Zebrafish lines, MO injection, and characterization of phenotypes

Tubingen strain (TU-AB) zebrafish were raised according to standard protocols as described earlier. Retinoblastoma (RB1) mutant lines were kindly provided by Dr. Indumathi Mariappan, BRIC inSTEM, Bangalore, India (ZFIN ID: ZDB-ALT-250929-16). All experiments were performed according to protocols approved by the Institutional Animal Ethics Committee of the Council of Scientific and Industrial Research, Centre for Cellular and Molecular Biology, India. Embryos were obtained from the natural spawning of adult fish, kept at 28.5°C, and staged according to hours after fertilization. ^55^ The endogenous *cep57* levels were depleted by using MO *cep57* translation blocker, 5’-AGGCTTTTGAGTTCGTCTCCATTAA-3′ and cep57 splice blocker, 5’-TGTCATAATGGACTCACAAGCTCAT-3’ (Gene Tools). For rescue experiments, cep57 mRNA or sox2 mRNA was *in vitro* transcribed (AM1340, Thermo Scientific), and 50pg of mRNA was co-injected along with c*ep57* splice MO in each embryo at the one-cell stage. The embryos were then analyzed for gross morphological defects and survival at later stages of development (till 5dpf) and were imaged in the Zeiss Stemi508 stereomicroscope.

### RNA isolation and Quantitative PCR

Total RNA was isolated from 100 embryos for each group using the RNA isolation kit (MN; 740955.50) as per the manufacturer’s protocol. cDNA was prepared using the PrimeScript 1^st^ strand cDNA synthesis kit (TaKaRa; 6110A). Quantitative real-time PCR (qPCR) was set up in the ViiA7 Real-Time PCR System (Applied Biosystems) using the prepared cDNA as a template along with PowerSYBR Green PCR master mix (Applied Biosystems, cat. no. 4367659). The following program was used: 40 cycles of 95 ^0^C -10min., 95 ^0^C -15 sec., 60 ^0^C -1 min., 95 ^0^C -15 sec., 60 ^0^C -1 min., 95 ^0^C -15 sec. Zebrafish β*-actin* was used as an internal control. The gene expression levels were measured in triplicate for control and morphants. The fold change of the genes tested was calculated using the 2^-ΔΔCt^ (the delta-delta-Ct) method. All primers used in this study are listed in Fig.S8.

### Antisense riboprobe preparation

For cloning sequence-specific exonic fragments to synthesize riboprobes, total RNA was isolated from a pool of 100 embryos (24hpf) of the TU-AB strain using an RNA isolation kit (MN; 740955.50). The total RNA was used to prepare cDNA using the PrimeScript 1^st^ strand cDNA synthesis kit (TaKaRa, Cat. # 6110A). Sequence-specific primers were used to amplify the specific gene fragment and cloned into the pGEMT easy vector, and the sequence was verified. The sequence-verified plasmid was linearised using Nco1, and antisense digoxigenin (DIG)-labeled riboprobes were synthesized using the DIG RNA labeling kit (Roche #11175025910). ^20^

### Whole Mount RNA in Situ Hybridization

The control and the morphant embryos of the desired developmental stage were dechorinated and fixed in 4% paraformaldehyde (PFA) in PBS overnight. The embryos were then washed with PBT (1× PBS and 0.1% Tween 20) and stored in methanol at -20 ^0^C overnight. Embryos were rehydrated using decreasing gradation of Methanol-PBT followed by pre-warmed hybridization wash buffer (50% formamide, 1.3 × SSC, 5 mM EDTA, 0.2%, Tween-20) and blocking in pre-warmed hybridization mix buffer (50% formamide, 1.3 × SSC, 5 mM EDTA, 0.2%, Tween-20, 50 μg/mL yeast t RNA, 100 μg/mL heparin) for 5 hours at 55 ^0^C. DIG-labelled RNA probe was added (1ng/ul) and kept overnight at 55 ^0^C. The next day, the embryos were washed with a prewarmed hybridization mix buffer followed by washes of a gradient of 2X SSC (20X SSC stock-3M NaCl, 0.3M sodium citrate, pH7.2), Hybridization wash buffer, 0.2XSSC and TBST (0.5 M NaCl, 0.1 M KCl, 0.1 M Tris, pH 7.5 and 0.1% Tween 20) at room temperature. The embryos were then given a TBST wash and then blocked in 10% heat-inactivated fetal bovine serum (Gibco; 16210-064) in TBST for 3 hours at room temperature. Anti-DIG antibody (1:5000) was added and incubated overnight at 4°C. The next day, four washes of TBST were given at room temperature, followed by four washes of NTMT (0.1 M NaCl, 0.1 M Tris-Cl at pH 9.5, 0.05 M MgCl2, 1% Tween-20) and stained with 1-step NBT/BCIP solution (Thermo Scientific, cat. no. 34042). Fresh NTMT was added to stop the reaction for both control and morphant embryos simultaneously and photographed using Zeiss Stemi508 stereomicroscope and Zeiss AxioZoom V16 microscope.^20^

### Western Blot analysis

Lysates of embryos younger than 1dpf were prepared by homogenizing approximately 100 embryos using RIPA buffer (20mM Tris–HCl, pH 7.4; 150mM NaCl; 5mM EDTA; 1% NP-40; 0.4% sodium deoxycholate; 0.1% SDS, and 1× Protease inhibitor cocktail). The lysis was carried out by incubating on ice for 15 minutes with intermittent vortexing. The homogenized lysates were centrifuged (12000 rpm for 20 min at 4°C), and supernatants were collected. The samples were then boiled in Laemmli buffer at 95^0^C for 10 mins. For 1dpf embryos, the embryos were first dechorinated and then deyolked by gentle pipetting using deyolking buffer (55mM NaCl, 1.8mM KCl, 1.25mM NaHCO_3_) followed by centrifugation at 3000rpm for 30secs. The embryos were then washed using wash buffer (110mM NaCl, 3.5mM KCl, 2.7mM CaCl_2,_ 10mM Tris HCl, pH8.5), centrifuged at low speed, and the supernatant was discarded. The pellet was then boiled in Laemmli buffer at 95^0^C for 10 mins. 20 μg of total protein was loaded in each lane and run on a 10% SDS-PAGE gel using Tris-Glycine SDS buffer for the detection of endogenous proteins. Proteins were then transferred to methanol-activated PVDF membranes (Merck Millipore, ISEQ00010) by electro-blotting. The membranes were then blocked either in 5% non-fat dry milk (sc-2324) or 5% BSA (Bovine serum albumin, for phosphoproteins) prepared in TBST (20 mM Tris HCl, pH 7.5, 150 mM NaCl, 0.05% Tween 20) for 1 hour. The blots were probed with primary antibody prepared in 1% blocking solution for 14-16 hours at 4^0^C. The blots were then washed with 1X TBST three times and incubated with HRP-conjugated secondary antibody for 1 hr. The blots were then washed again three times with TBST and developed for chemiluminescence signal using the chemiluminescent substrate (Thermo scientific-34577) and captured in the Vilber-Lourmat Chemiluminescence System. The primary and secondary antibodies used in this study are listed in Fig.S7.

### Nuclear cytoplasmic fractionation

Ice-cold 0.1% NP40 prepared in PBS was added to 200 dechorinated and deyolked, wild-type embryos and gently pipetted 5-10 times. 200 μL of lysate was collected as the whole embryonic lysate. The remaining sample was centrifuged at 2000 rpm for 5 minutes. The supernatant was carefully collected as the cytoplasmic fraction. To the pellet, ice-cold 0.1% NP0/PBS was added, and gentle pipetting was done. The sample was again centrifuged, and the supernatant was discarded. The pellet contains the nuclear fraction. 2X Laemmli buffer was added to all the three fractions, properly lysed, and heated at 95 °C for 10 mins. The purity of the fractions were confirmed by immunoblotting, probing with GAPDH (cytoplasmic marker) and LAMIN B1 (nuclear marker)^56^

### FACS-based whole embryo cell cycle analysis

Embryos at the desired developmental stage were fixed in 4% PFA/PBS and subsequently dechorionated and deyolked. To obtain a single cell suspension, PBS was added to the embryos and disintegrated using a 24-gauge syringe. The cell suspension was then centrifuged at 1000 rpm for 2 mins at 4 °C. The supernatant was collected in a fresh Eppendorf tube and centrifuged at 3000 rpm for 15 mins at 4 °C. The pellet was resuspended in DAPI (10 μg /mL) prepared in PBS. The single cell suspension was then centrifuged at 3000 rpm for 15 mins at 4 °C, resuspended in PBS, and analysed by flow cytometry. Cell cycle analysis was performed using Beckman Coulter Gallios flow cytometer and a total of 300000 singlet events were acquired for each group. The data was analysed using Kaluza Analysis 2.2 software, and plots were obtained with the histogram of the linear DAPI area using the J.V. Watson algorithm.

### Acridine orange staining

Acridine orange (AO) staining was performed to detect the apoptotic cells in 1dpf embryos. The control and morphant embryos were dechorinated, stained with 5mg/L AO for 1 hour in the dark, and then washed with embryo water. The embryos were anesthetized with tricaine, and the bright green dots in the embryos indicated apoptotic cells. The number of apoptotic cells in the head region was counted using Image J software. The apoptotic cells in the embryos were photographed by Zeiss AxioZoom V16 microscope.^20,57^

### TUNEL assay (terminal deoxynucleotidyl transferase dUTP nick end labelling)

Apoptotic cell death was assessed using the In Situ Cell Death Detection Kit, Fluorescein (Roche; 11 684 795 910), according to the manufacturer’s protocol with slight modifications. Manually dechorionated 1dpf control and morphant embryos were fixed in 4% PFA in PBS overnight at 4 °C, followed by methanol fixation for 48 hours. The embryos were then rehydrated using 50% methanol/PBS and washed thoroughly with PBS. Permeabilization was performed on ice for 15 min using 0.1% Triton X-100 in 0.1% sodium citrate and the embryos were incubated in the TUNEL reaction mixture containing 5 µL enzyme solution and 45 µL label solution for 2 hours 30 minutes at 37 °C in dark. The reaction was stopped by PBS washes and images were acquired using the Zeiss AxioZoom V16 microscope. Green fluorescent nuclei were considered TUNEL-positive, indicating DNA strand breaks in apoptotic cells.

### Genomic DNA isolation and RAPD PCR

Genomic DNA isolation was done for both control and morphant embryos using the Macherey-Nagel nucleospin tissue kit as per the manufacturer’s protocol. PCR reaction was set up using primer with sequence 5’-CCCGTCAGCA-3’ and the isolated DNA(200ng) as a template for 25cycles. The PCR products were run on 2% agarose gel and visualized under UV light. The difference in the PCR profiles of control and morphants forms the basis of the genomic template stability (GTS). GTS= (1-a/n)*100, where a is the number of polymorphic bands in morphants and n is the total number of bands in control. GTS is expressed as a percentage with control set to 100%. ^32^

### Comet assay (single cell gel electrophoresis)

Microscopic slides were pre-coated with 1% agarose by spreading a thin, uniform layer over the slide surface and allowed to solidify overnight at room temperature. 20 µL single cell suspension was mixed with 30 µL 0.8% low-melting agarose and carefully spread onto the agarose-coated slides. The suspension was covered with coverslips and allowed to solidify at room temperature. The cells were lysed in freshly prepared lysis buffer (2.5 M NaCl, 100 mM EDTA, 10 mM Tris-Cl pH-9.5, 1% Triton X-100, 10% DMSO) at 4 °C for 90 mins, followed by incubation in electrophoresis buffer (300 mM NaOH in 1mM EDTA) for 30 mins. Electrophoresis was carried out at 15 V and 400 mA for 15–20 mins at 4 °C. Subsequently, slides were neutralized with 0.1 M Tris-HCl (pH 7.5) for 15 mins (three washes), fixed in 100% methanol for 5 min, and rehydrated in Milli-Q water for 10 mins. DNA was stained with SYBR Green (1:50) for 10 mins in dark, washed with Milli-Q water, and visualized in the Zeiss AxioZoom V16 microscope. To assess DNA damage based on comet tail formation, the acquired images were quantified using ImageJ software. Comet tail length was measured for 200 individual nuclei over three biological experiments by tracing the extent of DNA migration from the head to the end of the tail. The mean tail length of morphant samples was compared to that of the corresponding control group, with increased tail length in morphants indicating elevated levels of DNA damage.

### Skeletal preparations - Alcian blue and Alizarin red staining

Five day-old control and morphants larvae were fixed in 4% PFA/PBS overnight at 4^0^C. The embryos were washed with 1XPBS and stored in methanol at -20 ^0^C overnight. The staining solutions needed for bone and cartilage staining were prepared: Solution A (0.04% Alcian blue, 125mM MgCl2, 70% ethanol) and Solution B (1.5% Alizarin Red S). 1 ml of solution A and 20ul of solution B were added to the larvae and kept overnight at room temperature in dark with constant rocking. The staining solution was removed, and the larvae were washed twice with water. Then, embryos were then transferred to bleach solution (1% KOH, 1.5% H_2_O_2_) for 1 hour at room temperature followed by a gradation of Glycerol-KOH washes. The embryos were then stored in 50% glycerol-0.25% KOH, and the larvae were imaged using Zeiss AxioZoom V16 microscope. For microcephaly analysis of the 5dpf larvae, the interpupillary distance (IPD) and the head size defined by the otic vesicle and the semicircle of eyes as posterior and lower boundary were quantified using Image J software.^58^

### Immunohistochemistry of zebrafish embryos

The control and morphant embryos at the 256-cell stage were fixed in 4% PFA/PBS at 4^0^C overnight, followed by methanol fixation. The methanol-fixed embryos were rehydrated with 50% MeOH/PBS followed by PBS washes and then blocked in 1% BSA/PBST (1X PBS and 0.1% Triton X-100) for 2 hours at room temperature. The embryos were then incubated with primary antibody, prepared in blocking solution overnight at 4 °C. PBST washes were given followed by incubation with appropriate secondary antibody overnight at 4 °C. The embryos were again washed with PBST, and DAPI (1µg/ml) was added for 10min, washed with 1X PBS, and stored at -20 °C until imaging. For micronuclei analysis, the single cells were first labelled with ActinRed (1:200, Invitrogen R37112) for 14 hours at 4 °C, prior to DAPI staining. The image acquisition was done using Leica TCS SP8 confocal microscope and Zeiss ELYRA7 with Lattice SIM2 Super Resolution Microscope. The mitotic phenotypes were quantified by 3D reconstruction of the confocal images using the IMARIS software. The primary and secondary antibodies used in this study are listed in Fig.S7.

### Immunoprecipitation and protein pull-down assay

Dechorionated and deyolked 1 dpf wild-type embryos were lysed in immunoprecipitation (IP) buffer (150 mM NaCl, 10 mM Tris HCl, pH 7.5, 1 mM EDTA, pH 8.0, 0.2 mM sodium orthovanadate, 1% NP40 and 1X Protease inhibitor) and centrifuged at 12,000 rpm for 20 mins at 4°C. The lysate was precleared by incubation with 25 µL of Protein A/G magnetic beads (Invitrogen: 78609) that was prewashed thrice with TBST (10 mM Tris HCl, pH 7.4, 15 mM NaCl, 0.01% Tween 20) at 4°C for 1 hour. Primary antibody (Cep57 and IgG) was added to the precleared cell lysate and kept at 4°C for 14-16 hours. 100 µL prewashed Protein A/G magnetic beads were added to the cell lysate and incubated overnight at 4°C. Using a magnetic stand, the flowthrough was collected and the beads were washed with IP buffer. For eluting the bound proteins, 2X Laemmli buffer was added to the beads, vortexed vigorously, and heated at 95°C for 10 mins. The eluate was then separated from the beads using a magnetic stand.

### Label-free quantification and proteome analysis

For comparing the changes in the total proteome upon gene knockdown, label-free quantification was done using mass spectrometry. Lysates of 24hpf control and *cep57* morphant embryos were prepared using RIPA buffer. 100ug of total protein of each sample was separated on 10% SDS-PAGE gel. The gel was stained using Coomassie blue and destained with acetic acid and methanol. The protein-loaded lane was excised from the gel and fragmented into multiple pieces. The gel pieces were then alternately washed with 50 mM ammonium bicarbonate and acetonitrile to clear off the Coomassie blue stain from the gel pieces. Further, the gel pieces were subjected to reduction reaction using 10mM dithiothreitol followed by alkylation using 55mM iodoacetamide. In-gel trypsin digestion was carried out at 37^0^C for 16 hours. The peptides were extracted using an extraction buffer (5% formic acid, 30% acetonitrile) and desalted using C18 reversed phase material-bound Zip Tips. The peptides were analyzed using the Thermo Scientific Easy-nLC 1200 system coupled to Q-Exactive mass spectrometer (Thermo Scientific). The peptides were resolved on a Thermo Scientific PepMap RSLC C18, 3μm, 100 Α, 75 μm x 15 cm column (ES900) using a 5% - 25% - 45% - 95% -95%-3% Solvent B gradient followed at 0.00, 35.00, 41.00, 46.00, 49.00, 53.00 and 60.00 minutes respectively, (Solvent A: 5% acetonitrile - 0.2% formic acid; Solvent B: 90% acetonitrile - 0.2% formic acid), in DDA mode (data dependent mode). The nLC eluent was sprayed into the QE with a spray voltage of 2.2Kv (kilovolts). The mass spectrometer was operated in the positive ion mode, and the settings with MS1 scans ranged over 400 – 1650 m/z, with AGC target, 3e^6^, resolution 70000. MS2 scans ranged over 200-2000m/z and resolution 17500. The data was acquired with the following set parameters: normalised collision energy 30, topN 10 peptides with charge exclusion 1, 6-8 and >8. The data was analysed using the Proteome Discoverer 2.2 software (Thermo Scientific), and the proteins were identified by performing a search against the *Danio rerio* Uniprot database (UP000000437). SEQUEST HT algorithm was used along with other analysis parameters such as trypsin enzyme specificity, carbamidomethyl static modification, mass tolerance set to 10ppm, and fragmentation tolerance set to 0.02 Da. Normalization mode was used for the total peptide amount along with pairwise ratio-based calculation. The percolator algorithm was used to validate peptide spectral matches (PSM), and a q-value cutoff of 0.01 (1% global False Discovery Rate, FDR) was used for proteome mapping, fold change, and GO analysis. ID mapping was performed using Uniprot, and Gene Ontology (GO) term analysis was performed using ShinyGO 0.76.2.^59,60^. Proteins with FDR<0.01 were qualified for quantification with a fold change cutoff ratio (FC) of >2.0 or <0.5 set as the threshold to identify differentially expressed proteins.

### Statistical analysis

Each experiment was repeated a minimum of three times independently. GraphPad Prism software was used for all statistical analyses. All the values are shown as mean with SEM unless specified. Data were analysed using an unpaired *t-test* for statistical significance. A *P* value less than 0.05 was considered statistically significant.

## Data availability

The mass spectrometry proteomics data have been deposited to the ProteomeXchange Consortium via the PRIDE [1] partner repository with the dataset identifier PXD062612.^61^

## CRediT authorship contribution statement

Conceptualization: MK, Methodology: SI and MK, Validation: SI and MK, Formal analysis: SI and MK, Investigation: SI, LPSM, AG and MK, Resources: MK, Data curation: LPSM, SI and MK, Writing – original draft: SI and MK, Writing – review and editing: MK, Visualization: SI and MK, Supervision: MK, Project administration: MK, Funding acquisition: MK

## Acknowledgments

We thank Dr. Santosh K. Guru, NIPER Hyderabad, for the Rb antibody and Dr. Indumathi Mariappan, BRIC inSTEM, Bangalore, for the Rb mutant zebrafish lines. We sincerely thank Mr. G. Srinivas, Advanced microscopy & Cell Sorting Facility, and Mr. B. Raman, Ms. Y. Kameshwari, and Mr. K. Ranjith Kumar, Proteomics Facility, CSIR-CCMB, for their technical assistance with the experiments. SI acknowledges CSIR, India, for the research fellowship. MK thanks the Anusandhan National Research Foundation (ANRF), Govt. of India (ANRF/ECRG/2024/000070/LS) and Council of Scientific and Industrial Research - Centre for Cellular and Molecular Biology (CSIR-CCMB), Govt. of India, for supporting and funding this research.

---

**Figure S1:**

A: Sum projection confocal images of 256 cell stage WT embryos showing centrosomal staining by γ tubulin (red), nuclear staining of Cep57 (green, white asterisks), and DNA (blue/DAPI). Scale bars, 50µm. B: RT-PCR analysis of *cep57* transcripts in different developmental stages. β-actin was used as the loading control. C: Western blot showing Cep57 levels in different developmental stages till 24hpf. Gapdh was used as the loading control. D: Whole-mount RNA *in situ* hybridization showing *cep57* expression during early development. Scale bars,100µm.

**Figure S2:**

A: Quantification showing percent phenotype of control and *cep57* morphants injected with 1mM *cep57* translation blocker morpholino. Data are shown as mean ± SEM, *p<0.05, **p<0.01, ***p<0.001. n=3 for each experiment. B: Quantification showing percent phenotype of control and *cep57* morphants injected with 0.1mM and 0.5mM *cep57* translation blocker morpholinos. Data are shown as mean ± SEM, *p<0.05, **p<0.01, ***p<0.001. n=3 for each experiment. C: PCR-based validation of intron retention in *cep57* morphants injected with splice morpholino and rescue of splicing event when coinjected with *in vitro* transcribed cep57mRNA. mRNA alone-injected embryos showed correct splicing. No template control (NTC) was used as a negative control. D: Gross morphological analysis of *cep57* morphants P0, P1, P2, and P3 at different stages of larval development.

**Figure S3:**

Quantitative real-time PCR plots showing relative transcript levels of microcephaly markers *mcph1*, *wdr62*, *ankle2*, *map11*, *kif14,* and *aspm*. All data are shown as mean ± SEM. *p<0.05, **p<0.01, ***p<0.001. n=3 for each experiment.

**Figure S4:**

Spatiotemporal gene expression analysis by *in situ* hybridization in control and *cep57* morphants at 6 somite and 24hpf stage. A, B, C: *Crestin, snail2,* and *sox10* expression in the 6 somite control and *cep57* (red asterisk) morphants. D, E, F: *Crestin, snail2,* and *sox10* expression in 24hpf control and *cep57* (double red asterisk) morphants. G: *Sox2* expression in 24hpf control and *cep57* (red asterisk) morphants. Scale bars, 100µm. H: *Tbxta* expression in 24hpf control and *cep57* (red arrow) morphants. Scale bar, 50µm.

**Figure S5:**

A: PCR-based RAPD assay showing polymorphic bands (red arrows) in control and *cep57* morphants. B: Quantification of RAPD assay to estimate genomic instability in control and *cep57* morphants. All data are shown as mean ± SEM. *p<0.05, **p<0.01, ***p<0.001. n=3 for each experiment. C: Sum projection confocal images of 256 cell stage control and *cep57* morphant embryos showing mitotic cells by PH3 (green) and DNA (blue/DAPI). Scale bars, 50µm. D: Western blot showing Cyclin E levels in in control and *cep57* morphants. Gapdh was used as the loading control.

**Figure S6:**

GO analysis of all the upregulated and downregulated proteins in *cep57* morphants, respectively.

**Figure S7:**

List of all the antibodies used in this study.

**Figure S8:**

List of all the primers used in this study.

**Figure S9:**

List of all differentially expressed protein from LC-MS/MS analysis.

