## Supplementary figures and images for "Cep57 is a cohesin-associated regulator of chromosome segregation, cell cycle progression and genome stability in early embryos"

### supplementary_figs

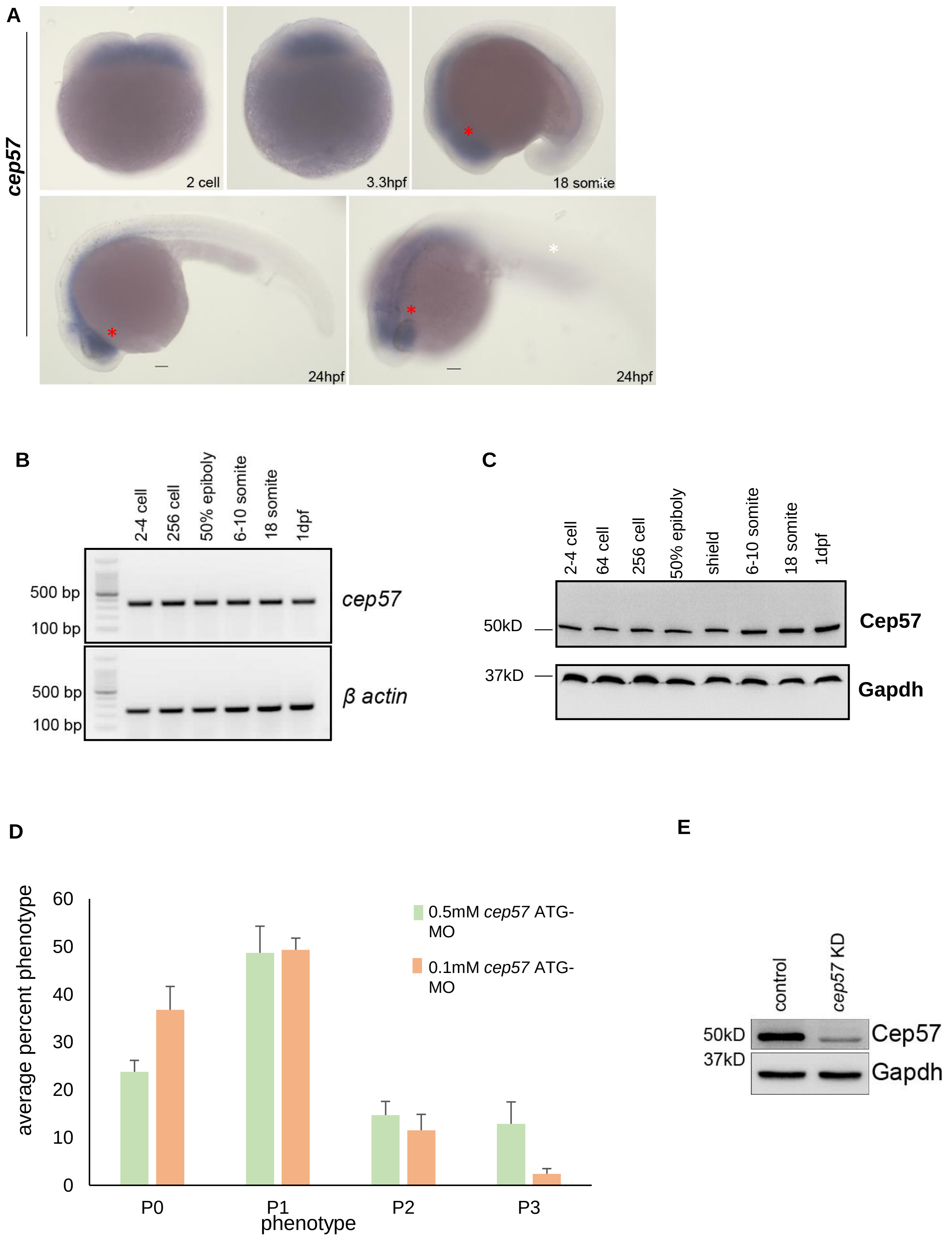

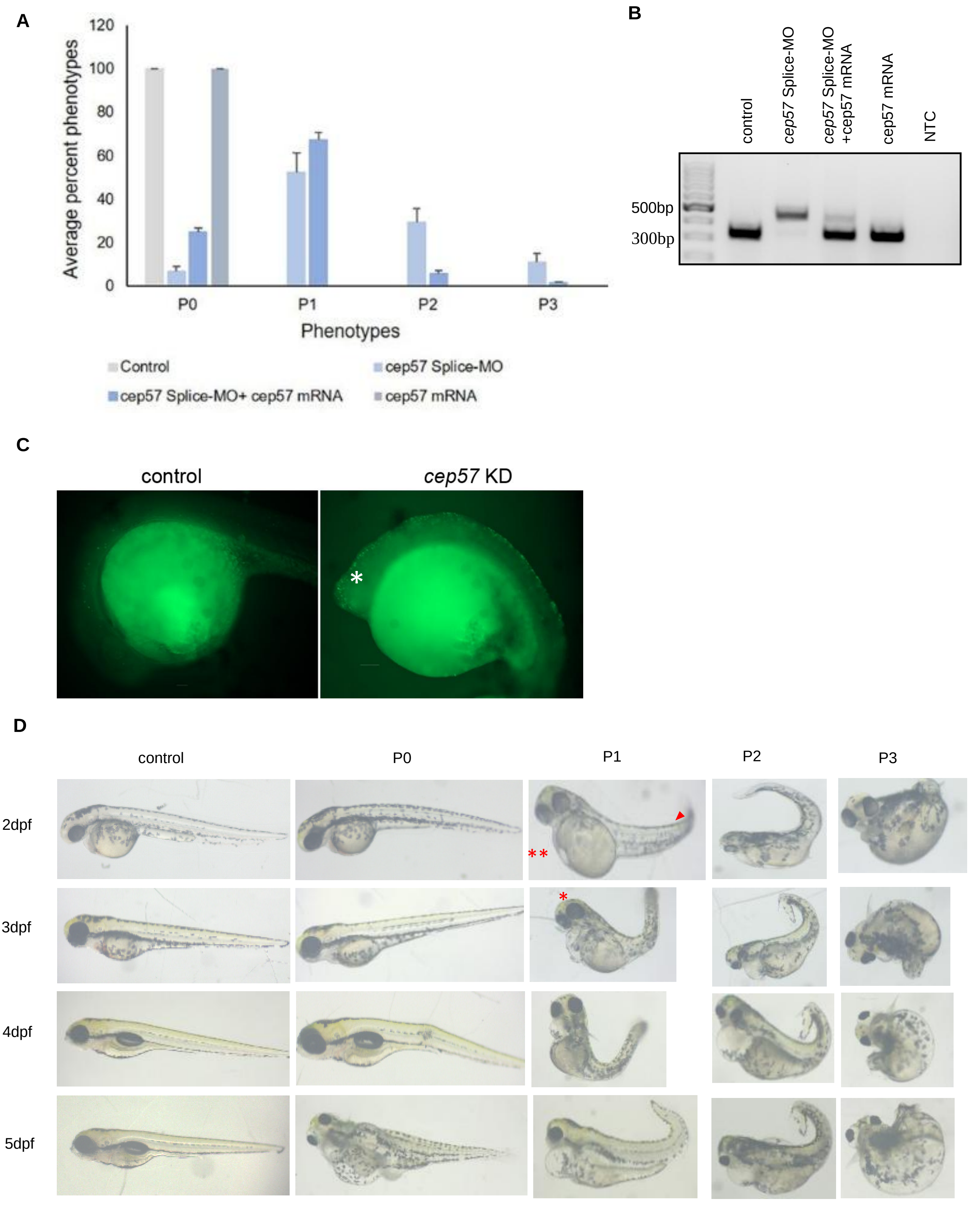

#### Figure S3

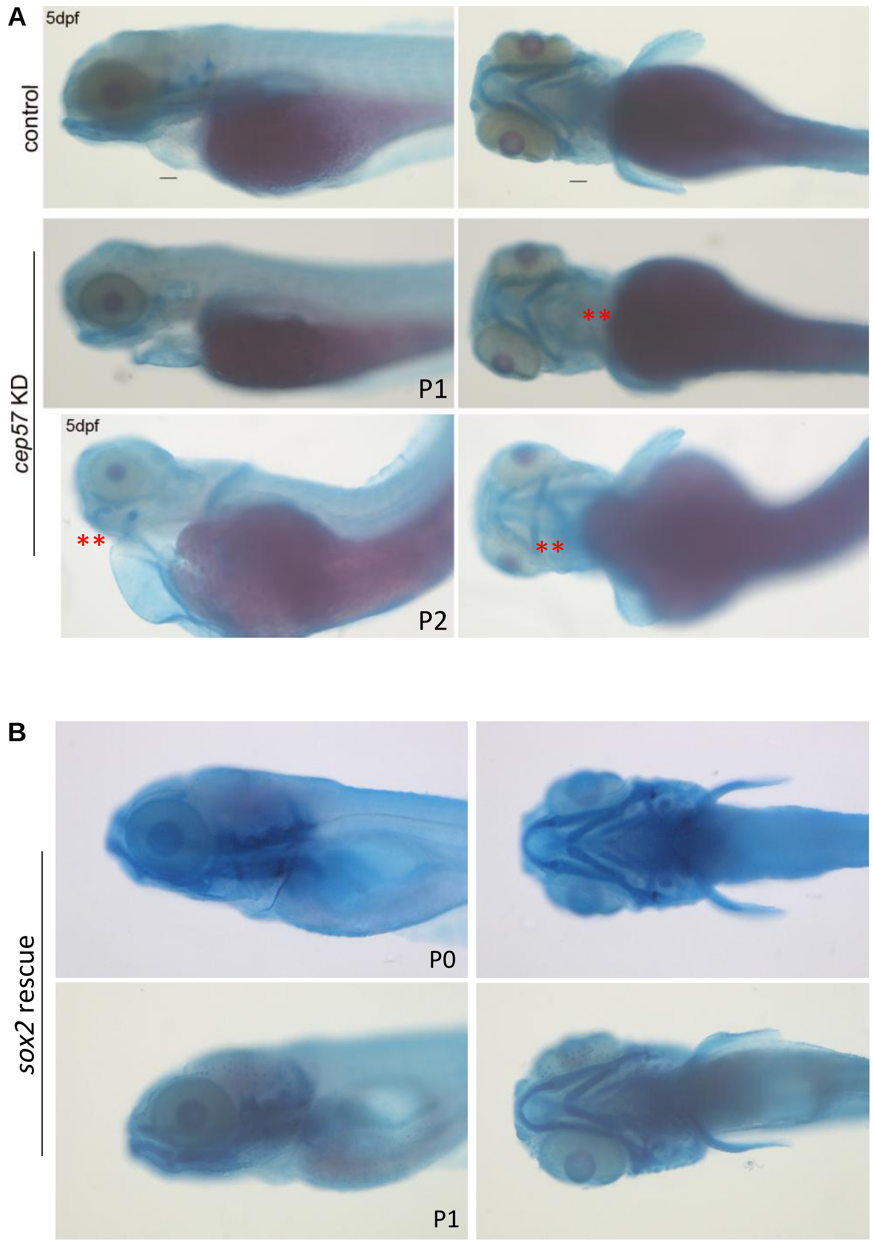

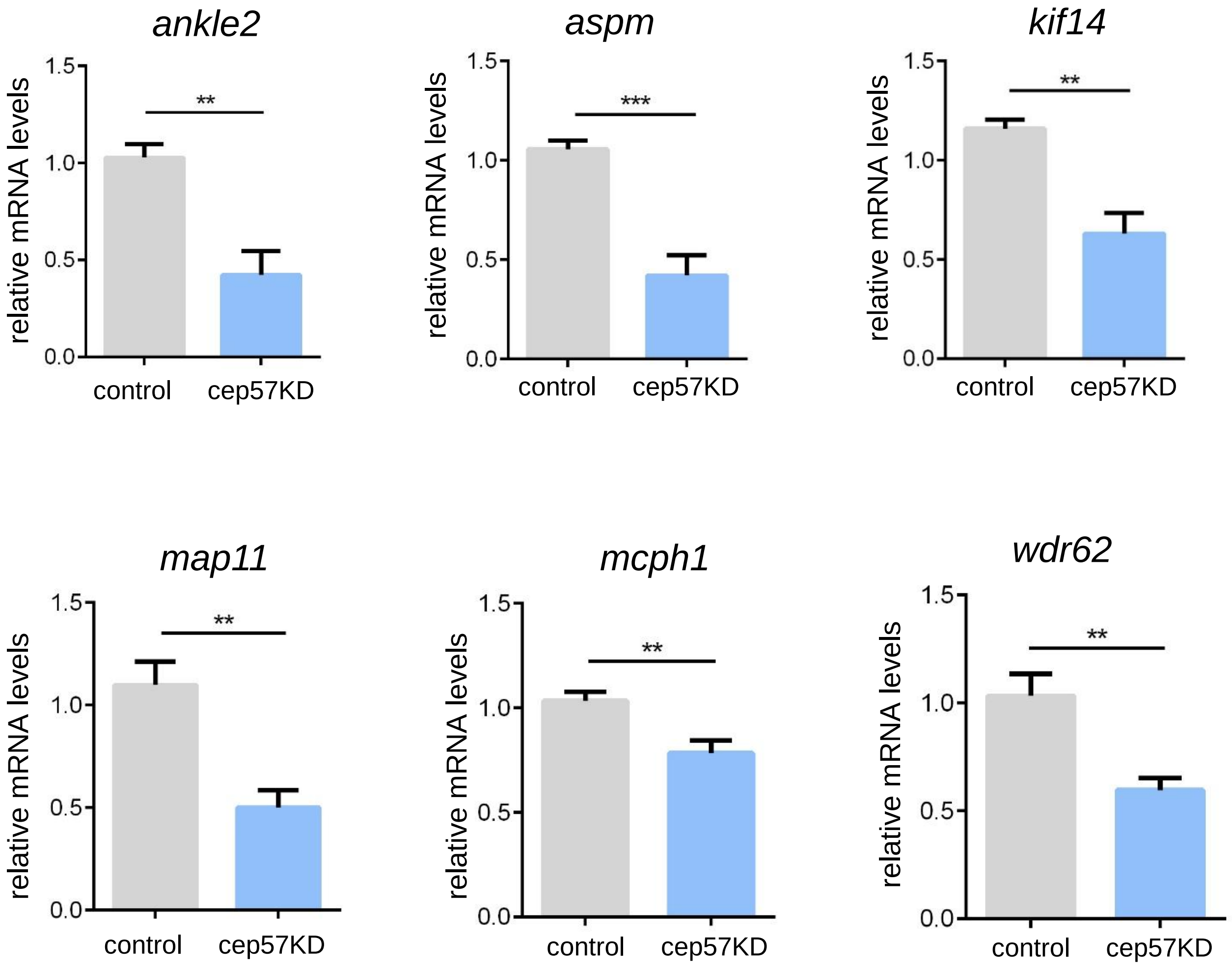

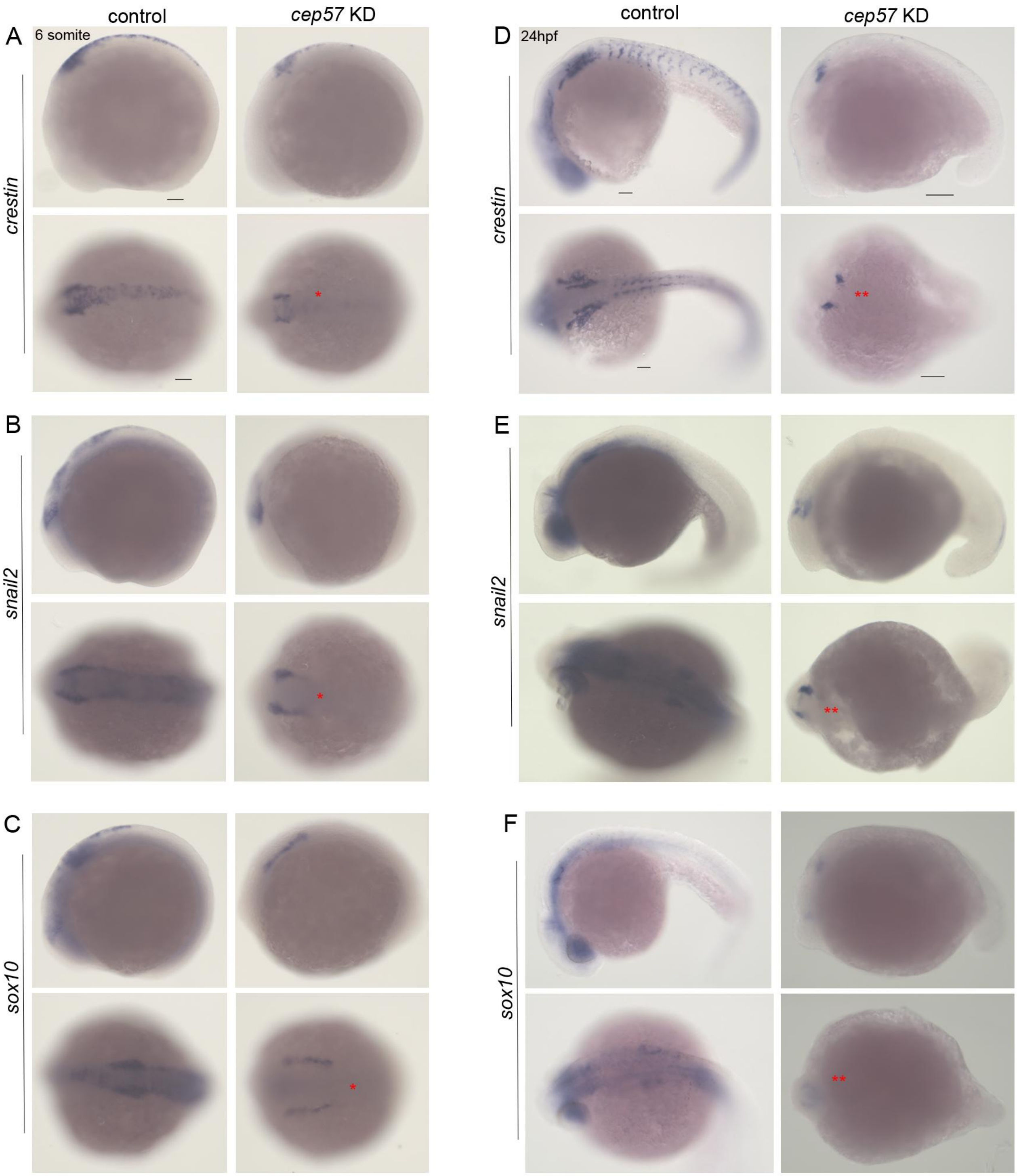

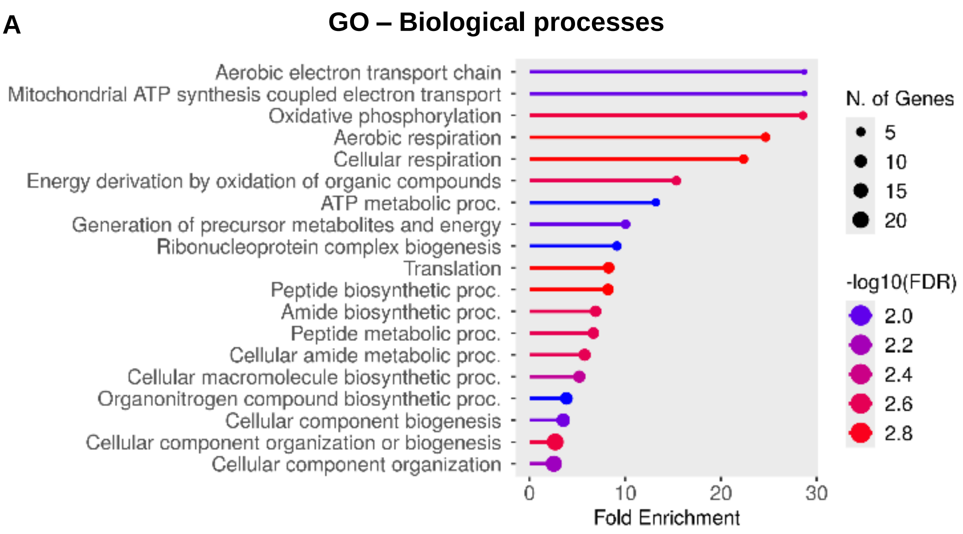

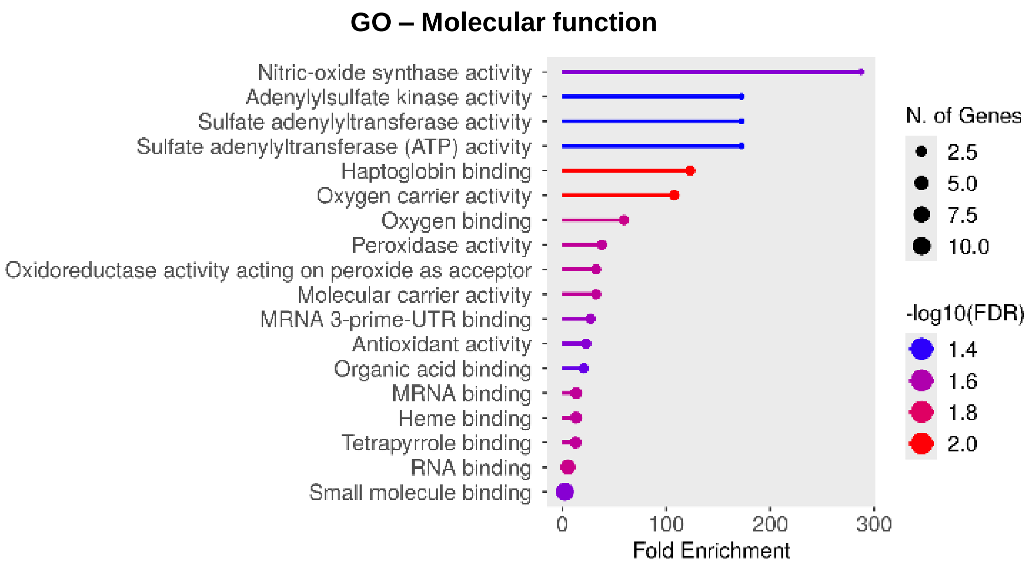

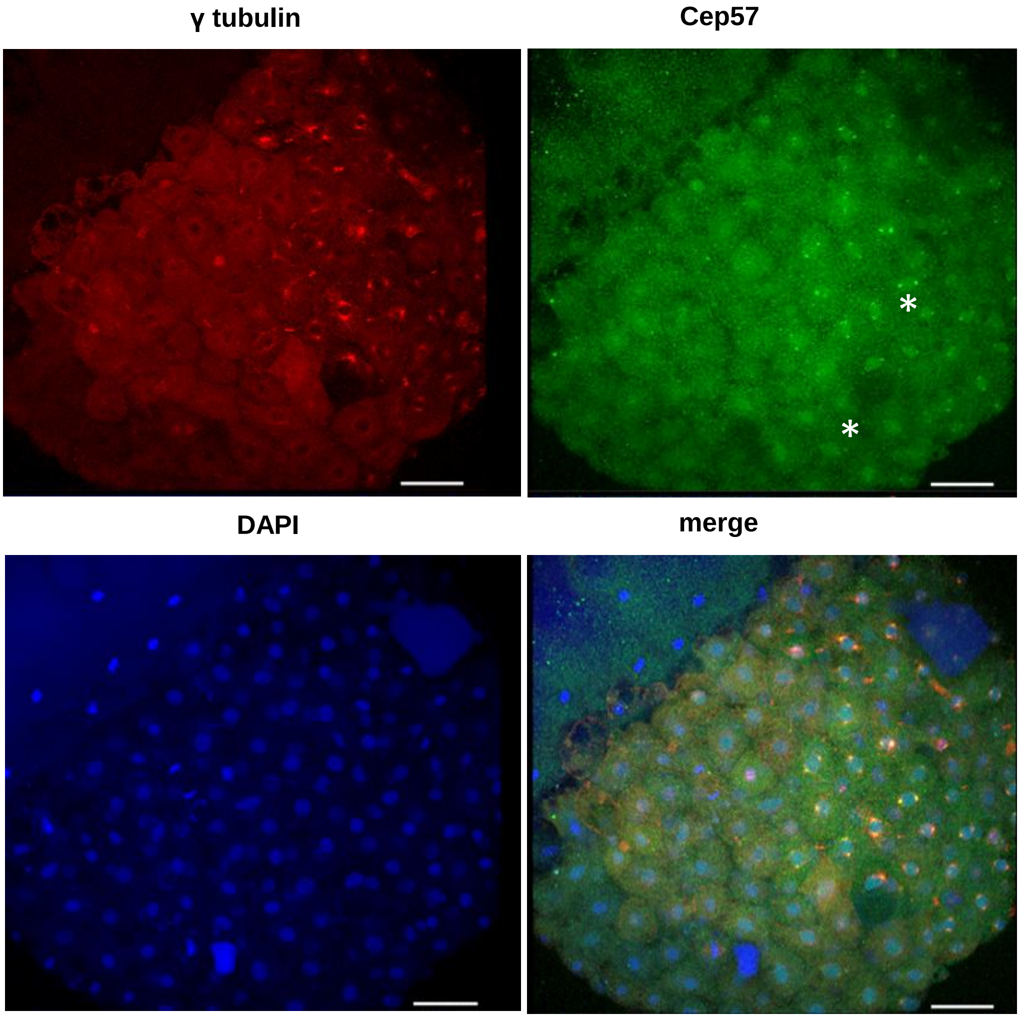

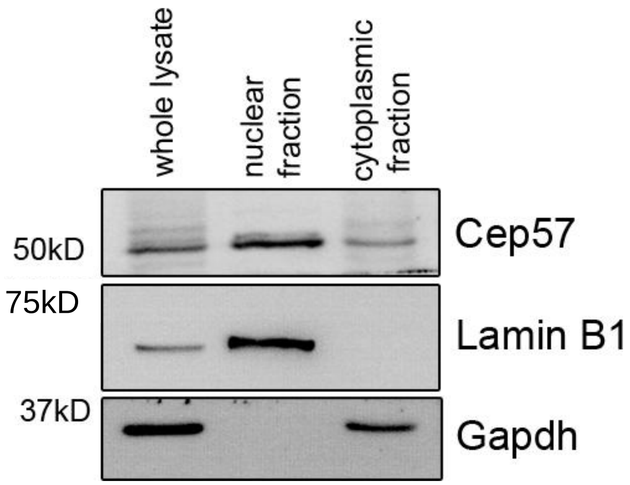

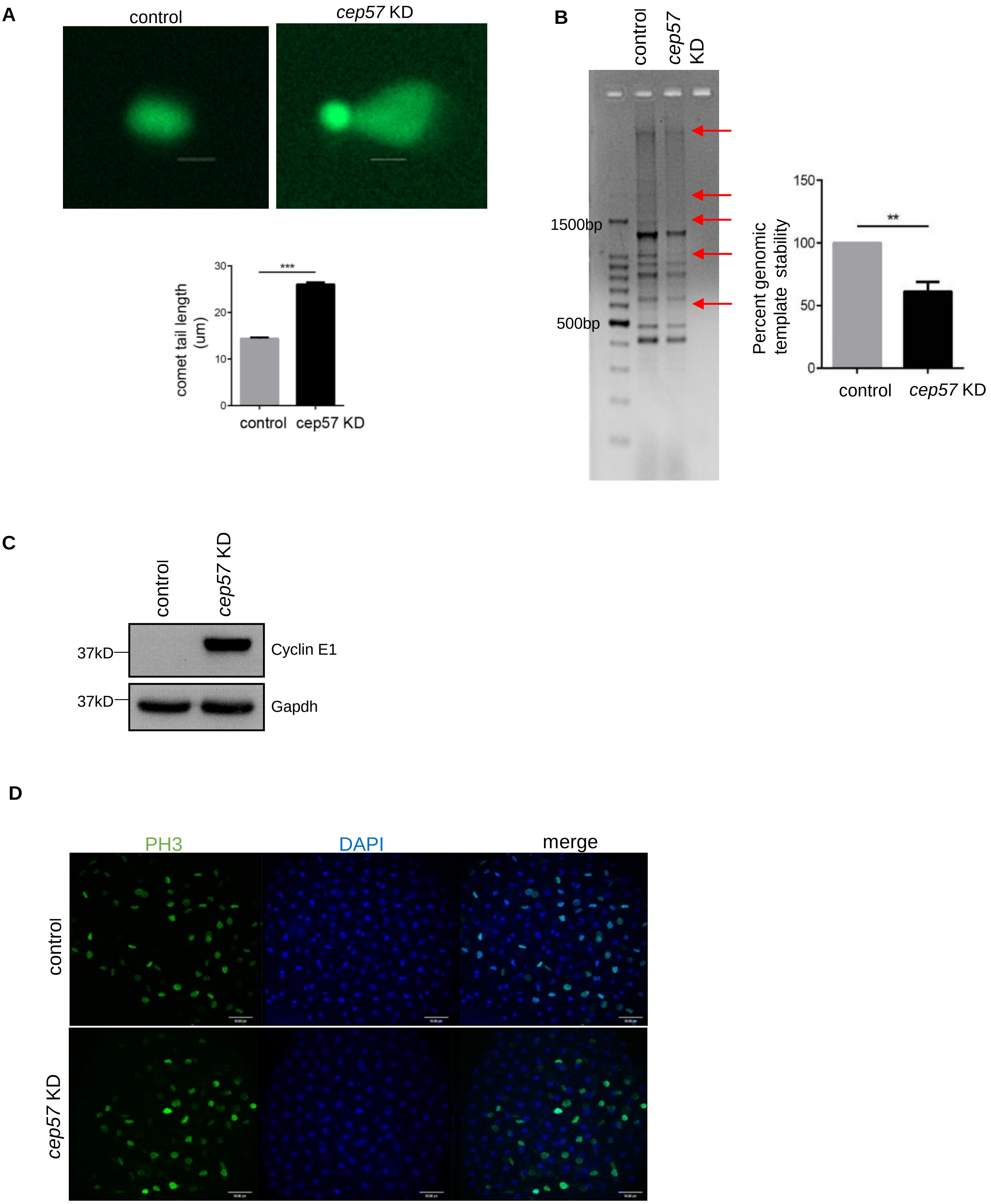
